# Reducing indoor particle exposure using mobile air purifiers - experimental and numerical analysis

**DOI:** 10.1101/2021.07.23.453308

**Authors:** Adrian Tobisch, Lukas Springsklee, Lisa-Franziska Schäfer, Nico Sussmann, Martin J. Lehmann, Frederik Weis, Raoul Zöllner, Jennifer Niessner

## Abstract

Aerosol particles are one of the main routes of transmission of COVID-19. Mobile air purifiers are used to reduce the risk of infection indoors. We focus on an air purifier which generates a defined volumetric air flow through a highly efficient filter material. We investigate the transport of aerosol particles from an infected dummy equipped with an aerosol generator to receiving thermal dummies. For analysis, we use up to 12 optical particle counters to monitor the particle concentration with high spatial resolution. Based on the measurement data, a computational fluid dynamics (CFD) model is set up and validated. The experimental and numerical methods are used to investigate how the risk of infection suggested by the particle exposure in an exemplary lecture hall can be reduced by a clever choice of orientation of the air purifier. The particle concentration at head height deviates by 13 % for variations of location and orientation. Finally, CFD simulation was used to monitor the particle fates. The steady simulation results fit quite well to the experimental findings and provide additional information about particle path and for assessing comfort level due to air flow.

**Practical implications:** Different installation locations and operating conditions of the air purifier are evaluated and the use of thermal dummies mimics the conditions of practical use cases. The measurement results show the integral particle mass over time in the “faces of the dummies”, representing the potentially inhaled particle load of persons present in the room. At an air change per hour of 5, the cumulated PM1 mass at head level was reduced by 75 %, independently of the location of the infected dummy, compared to the “natural decay” case showing that filtration is an effective means of reducing aerosol particle concentrations. It turns out that obstructing the outlet stream of the air purifier may be particularly advantageous.

## I). Introduction

The COVID-19 pandemic has implied large restrictions to public and private life and has far-reaching effects on society, culture, science, and the economy. It is well-known that a major route of infection with SARS-CoV-2 is the transmission by aerosol-borne viral pathogens [1]. These virus-laden aerosols may be emitted when talking, shouting, singing, coughing, sneezing or simply breathing. Small aerosol particles may remain suspended in the air for hours [2]. Infections may occur by proximity, when aerosols emitted by one person are directly transported towards another person. On top of this direct infection route via droplets an indirect infection route exists indoors, where aerosol particle concentration, and thus, infection risk, increases with time depending on the number of persons present, their activity, and the air volume within the room [3]. Since people spend over 90 % of their time indoors and several persons may be infected at a time, indoor situations are most crucial for SARS-CoV-2 transmission [4]. Personal protective measures, such as masks will never be able to remove all particles, therefore ventilation is of utmost importance as an additional measure. While opening windows very regularly (if possible) is helpful under warm conditions, air filtration is an important supplement in spring, autumn and winter. Discussions about ventilation strategies by opening windows, the use of face masks during class, lectures, or office work, and the possible risk of infection are numerous. Understanding and controlling aerosols seems to be the key mechanism to minimize the infection risks. In this context, computational fluid dynamics (CFD) modeling is a powerful tool to investigate aerosol particle transport and fate. Also, recent advances in sensor network technology provide the possibility to measure particle concentrations not only at few locations within a room, but to monitor aerosol particle concentration as a measure of infections risk at the locations where particles are potentially inhaled.

While the infection risk through air borne transmission is dependent on multiple factors, mainly the number of emitted virus-laden particles, the half-life of the virus as well as the number of inhaled viable viruses, lowering the concentration of virus-laden particles in the air is key to lowering the infection risk of occupants via the indirect infection route indoors [5]. This can be achieved by either diluting the air with virus-free fresh air or by removing the aerosol particles using highly efficient filters. While critical places such as operating rooms achieve this through Heating, Ventilation and Air Conditioning (HVAC) systems with air changes per hour (ACH) ranging from 15 to up 40 [6], most HVAC systems in classrooms, theaters and offices are not designed to reduce the infection risk, but rather to keep CO_2_ concentration below the recommended level of 1000 ppm [7].

Mobile air purifiers represent a chance to reduce aerosol particle concentration by removing particles from the air using highly efficient filters. For this purpose, ACH of 5 to 6 are recommended [6, 8]. Multiple studies have investigated the decay rates of aerosol particle concentration using air purifiers at varied ACH. Burgmann et al.[9] showed that at an ACH of 6 the aerosol particle concentration is reduced by approximately 80 % within 30 minutes. Kähler et al. have reported decay rates ranging from 1.5 to 3.8 resulting in half-life times between 10 min and 27 min depending on the ACH. In order to minimize the dwell time of the aerosol particles, Kähler et al. recommend placing the air purifier in the center of the room if possible [8]. Curtius et. al. were able to report a 95 % reduction of aerosol particle concentration after 37 min using multiple purifiers to obtain the recommended ACH of > 5. The decay rate determined at an ACH of 5.7 was (0.107± 0.01) min^−1^, while the natural decay rate was (0.020 ± 0.01) min^−1^ [10].

Since the air velocity is larger at the outlet than at the intake, the purified air is discharged at greater distance [11]. On account that aerosol particles move with bulk air [12], this presents a chance for the air purifier to disperse the aerosol particles throughout the room, rather than removing them. [13] showed that the clean air delivery rate (CADR) of an air purifier in a small room of 70 m^3^ is by and large independent of its position. By placing the air purifier in a particular disadvantages position the decay rate of the particle concentration and therefore the CADR decreased [13].

The CADR is determind from Equation (1) by measuring the reduction rate *k_purifier_* of aerosol particles (particle size ranging from 0,09 μm to 11 μm) taking the natural decay rate *k_natural_* into account in a standardized test chamber *V_room_* [14], while the ACH is calculated by the volumetric flow rate of the air purifier divided by the room volume. This entails that although the ACH of a particular air purifier is in line with recommendations the CADR can be greatly reduced by a disadvantageous installation.

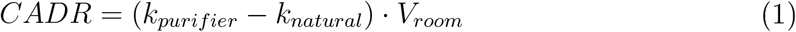

Investigation of the effectiveness of air purifiers in reducing indoor aerosol particle concentration often combine experimental and numerical methods. This allows for the examination of multiple cases using validated CFD models while minimizing experimental effort. Multiple studies have modeled the transport of aerosol particles indoors. In these publications various results of the importance of humidity and the thermal effects to a numerical particle simulation are discussed.

Feng et al. [1] showed that the most numerical studies report that condensation and evaporation due to humidity have a negligible effect on the particle distribution. Chen et al. [15] and Farkas et al. [16] focus on the deposition of multicomponent droplets and evaporation in a human respiratory tract. Feng et al. [1] faces this question of humidity and evaporation effects in indoor conditions, like in this study of a lecture hall. The evaporation time of water aerosols with a diameter of a few micrometers has to be shown to be less than one second [17]. Mutuku et al. [18] investigates different turbulence models and solver algorithms for particle simulations. In the most studies, an Euler-Lagrange method is used to simulate a multiphase particle flow [19]. Due to the low mass loading, a one-way coupling is used for fluid particle interactions [20]. Regarding the continuous phase, it can be assumed that an indoor air flow is incompressible and turbulent [9]. For turbulence modeling, Abuhegazy et al. [20] uses a *k* – *ε* model following the Reynolds-averaged Navier-Stokes equations (RANS) approach, since time-averaged results are often of interest. Modeling the discrete phase, the particle diameter distribution is important for the simulation [18]. Brownian particular motion can be neglected because the particle diameters are still too large [20]. In the simulation of air purifiers using highly efficient HEPA filters, it can be assumed that 100 % of the particles get removed [9], In contrast to this study in a lecture hall, Dbouk et al. [19] investigated aerosol dispersion in very confined spaces, as in an elevator and pointed out that the location of the inlets and outlets have a significant influence on the aerosol distribution. A larger outdoor environment is investigated by Gorbunov et al. [21] who shows in a simulation, that aerosol particles can travel over 30 m distance. Bathula et al. [22] uses simulation to investigate how long infectious particles remain in a room. This has practical implication regarding the safety of medical staff. Pyankov et al. [23] presents a study of time-dependent inactivation of MERS-CoV in ambient air under climatic conditions representing a common office environment. An alternative way of reducing infection risk is to inactivate the virus rather than removing aerosol particles. Rezaei et al. [24] investigates virus elimination by heat in air conditioning systems, to reduce the amount of contaminated particles. Another study on virus inactivation by UV-C irradiation is investigated by using numerical simulations of virus-laden droplets [25]. However, particles are still contaminated on the path from the source to the air cleaner, like when using common filtrating air purifiers.

The literature review shows investigations of the general dispersion of aerosols in different rooms and under different conditions. However, the studies do not focus on the particle load that the persons are exposed to at their seat positions. Therefore, in this study we investigate the particle concentration at positions where particles may be inhaled by persons using a high spatial resolution.

Therefore, the purpose of this work is to

- assess the influence of the position and the orientation of an air purifier on the aerosol particle concentration within the room and to identify preferable cases.
- investigate particle exposition of persons by measuring particle concentrations using a sensor network consisting of 12 PM1 sensors exactly at the locations where particles may be inhaled.
- include the influence of thermal buoyancy by mimicking the effect of persons present in the room using, thermal dummies”.
- investigate particle concentrations for the case that a sink (air purifier) fights against a source (infected person) and by studying commonly considered decay.
- validate a CFD model based on the measurements in order to 1) increase process understanding by allowing for a visualization of the flow field in a room and 2) allow for the investigation of situations which can hardly or not be considered experimentally.

In Section 2, we give an overview on the situation in the lecture hall considered, the experimental material, and the setup. Next, in Section 3, we introduce the mathematical and numerical model and give an overview on the considered cases. The numerical model is validated in Section 4 and both numerical and experimental results are presented and discussed. Finally, we sum up and give an outlook on future research in Section 5.

## II). Situation and experimental setup

Experiments were performed in a lecture hall at Heilbronn University of Applied Sciences. The dimensions of the room are 11 m × 8,5 m × 3 m. Due to the ascending rows, the ceiling height in the back of the room is significantly lower, resulting in a room volume of approximately 250 m^3^. The room provides seating for up to 80 students plus 1 professor. The total window area is 20 m^2^, of which 6 m^2^ can be used for ventilation.

The HVAC system is turned off during the entirety of the experiments. The room is equipped with a portable air purifier capable of a maximum volume air flow of 2500 m^3^/h, delivering a maximum ACH of 10. The air purifier is fitted with an HEPA H14 filter which removes at minimum 99,995 % of particles at the most penetrating particle size (MPPS) [26]. The inlet is located at the front side, the outlet perpendicular to one side of the device. Up to nine thermal dummies are deployed to simulate the heat flux contribution of persons in the room, in addition to serving as obstacles for the flow. The dummies are made of cardboard equipped with lightbulbs emitting approximately 75 W of thermal energy in accordance with the thermal contribution of students in a lecture scenario specified by DIN EN 16798-1 [7].

The room is equipped with up to 12 optical particle counters (OPCs), made up of four high-end devices (Fidas Frog, PALAS) and a network of 8 low-cost OPCs (SPS 30, SEN-SIRION). The temperature, ambient pressure, and relative humidity are measured at each point. Each device measures aerosol particles in the size range of 0,18 μm to 18 μm (Frog) or 0,3 μm to 10 μm (SPS 30). The high-end devices are deployed to measure particles at table height, while the low-cost sensors are placed at ceiling height as well as at “face height” of the dummies. The aerosol particles are produced using an atomizer (AGK 2000, PALAS) with a sodium chloride solution. The emitted particle size distribution is adjusted via the mass concentration of the saline solution, while the mass flow is set through the applied compressed air pressure.

### Influence of position, orientation and flow rate

In the first experimental set-up the influence of position, orientation and flow rate of the air purifier on the spreading and degradation of aerosol particle concentration is investigated. For this purpose, the aerosol particle are emitted in a fixed place in the center of the room using compressed air at 3,5 bars and a sodium chloride solution of 2.5 wt-%. The particle counters are positioned at table height at the measurement points MP 26, MP 58, MP 86 and MP 52 (see Figure 1). The experiments consist of two phases. In the first phase, the aerosol particles are emitted for a duration of 20 minutes, after which the atomizer is shut off and the decay of the particle concentration is monitored. The air purifier is positioned in four different locations throughout the room and at each position the orientation is varied so that inlet and outlet point in different directions into the room or against an obstacle, in this case either the window front or the wall. The following positions are identified as viable places to set-up the mobile air purifier (see Figure 1):

- NE - North-East near window panel
- N - North near window panel
- NW - North-West near window panel
- SW - South-West near the wall

**Figure 1:**
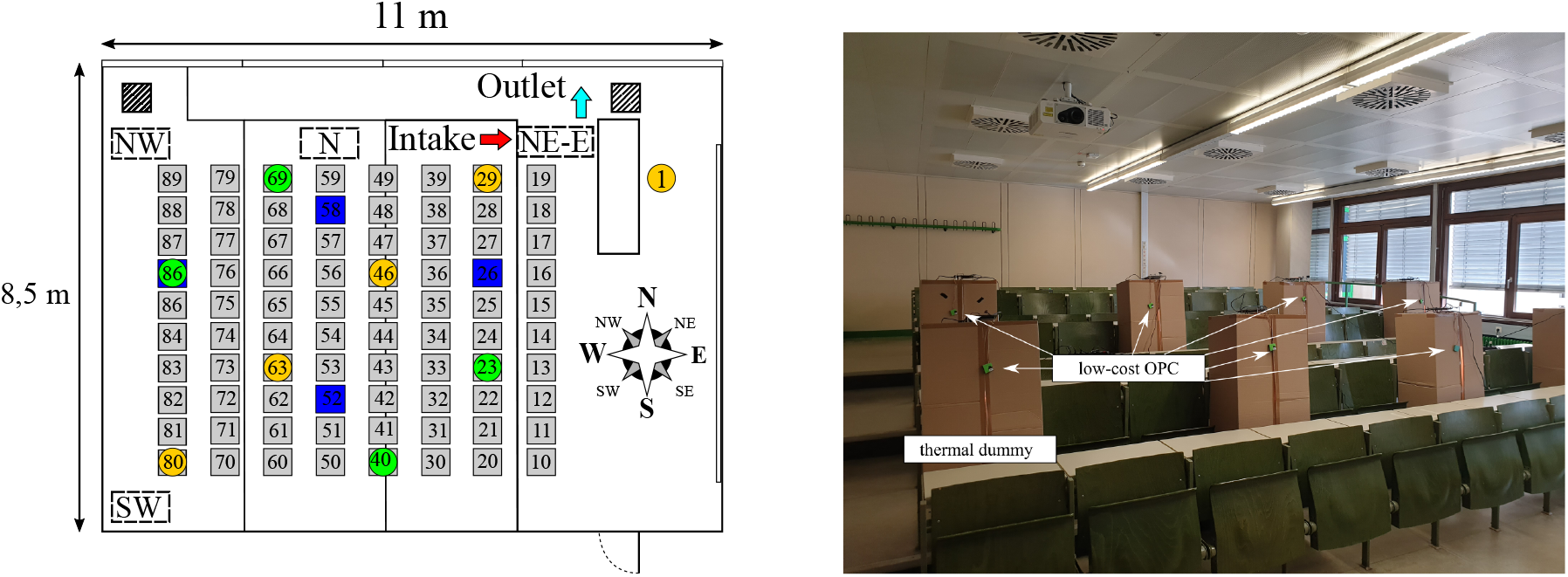
left-hand side: position of the air purifier, the receiving and emitting thermal dummy (marked by green and orange circles) and the high-end OPCs (marked blue); right-hand side: picture of the lecture hall with thermal dummies and sensor network

At most locations, particular orientations are disqualified as viable set-up options due to proximity to students or the professor and therefore likely exceeding air velocity limits for thermal comfort. The position and orientation of the air purifier are encoded using the cardinal direction relative to the compass seen in Figure 1 to identify the location, appended by the air flow direction at the intake of the air purifier. This leaves the following cases for the experimental set-up:

- NE-N; NE-E
- N-E
- NW-N; NW-W; NW-S; NW-E
- SW-N; SW-W; SW-S

The effectiveness of the operation parameters of the mobile air purifier (position, orientation) is evaluated by numerical integration of the mass concentration of particles smaller than 1 μm (PM1 fraction) over the duration of the experiment. This is due to fact that virus-laden droplet nuclei are known to be in a size regime < 5 μm. The value is then multiplied by 1.333 · 10^−4^ m^3^/s, representing the volume flow rate of a breathing person in a relaxed condition. This gives an average mass of PM1. which is inhaled at the measuring point over the duration of the experiment. Split up into the charging and decaying phase of the experiments, this value is used to identify the PM1 load at each measurement point throughout the classroom. By fitting the decay curve with an exponential decay function

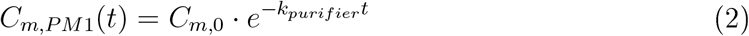

the decay rate *k_purifier_* at each measurement point is determined.

### Influence of particle source on local particle exposure

The second experimental set-up focuses on the influence of the position of the aerosol particle source on the distribution and decay of particles for fixed positions and orientation of the air purifier. Derived from the first experimental set-up, the following three cases are identified for further investigation: SW-W, SW-N, SW-S. Theses cases repesent a corner installation with an obstructed outlet (SW-W), obstructed outlet and intake (SW- N) and unobstructed outlet and intake (SW-S). The fourth case is a reference case where the air purifier is turned off. Nine thermal dummies are positioned throughout the room at maximum distance to each other, as shown in Figure 1. The location of the aerosol particles source is varied. The aerosol particles were emitted at seats 29, 46, 63 and 80. The particle concentration at table height as well as “face-height” of the dummies is measured.

## III). Numerical setup

In addition to the experimental studies numerical flow simulations are performed to achieve a deeper insight into the indoor air flow and the aerosol distribution. Additional cases are considered that can hardly be implemented in the experimental setup.

### Numerical approach

Two different numerical approaches of multiphase flows have been established. The Euler-Lagrange approach and the Euler-Euler approach. In particle laden flows with a low mass loading the Lagrangian approach is advantageous. It is assumed, that a mass point approach with approximation of the forces by particle volume is sufficient. Therefore, the particles are assumed to be spherical. Due to the low mass loading of less than 10 %, an Euler-Lagrange approach with a discrete phase model (DPM) and a one-way-coupling is used for the numerical simulations. The continuous phase influences the discrete phase by friction and turbulence. However, the particles do not influence the flow [27].

### Govering equations for the Euler-Lagrange approach

#### Continuous phase

In this case the flow field is calculated before the discrete phase calculation. Therefore, there is no influence of the particles on the flow field and the classical Navier Stokes equations are solved [28].

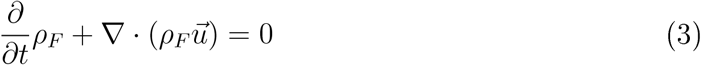

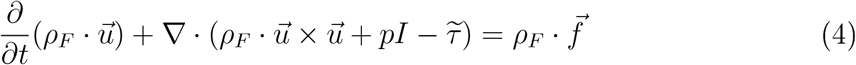

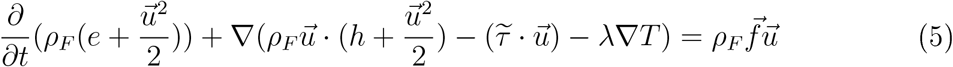

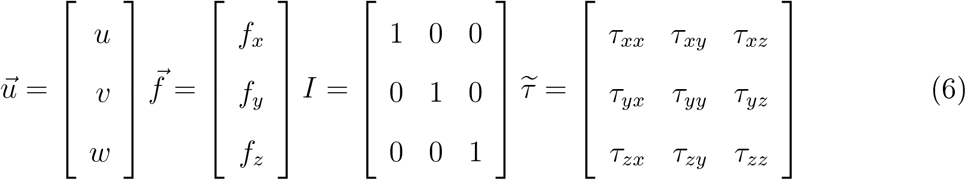

Index *F* represents fluid parameters. The density is denoted by *ρ* and the flow velocity vector 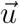, the pressure *p*, shear stress tensor 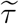. Additional accelerations like the gravitation is considered in the variable 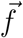, the specific internal energy *e*, specific enthalpy *h* and the Fourier’s law of heat conduction –λ∇*T*. Furthermore, the RANS averaging and the transport equations of the *k* – *ε* turbulence model and other additional equations like the incompressible ideal gas law are considered.

#### Discrete phase

Because an exchange of mass, momentum, and energy is not considered, the Lagrangrian equations can be highly simplified. The particle inertia can be written as in [29].

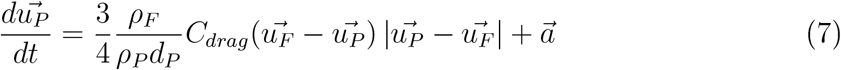

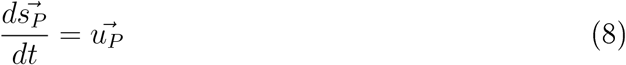

with diameter *d* and the drag coefficient *C_drag_*, which is calculated by the spherical drag law. Index *P* represents particle parameters. All other additional accelerations like the Saffman liftforce and gravitation in this case are represented by 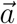. The vector 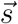 represents the position of the particle.

### Discrete phase model

In this parameter study, a steady particle tracking based on a fixed steady airflow is solved with ANSYS Fluent. Due to the low mass fraction, the one-way-coupling is applied. Nevertheless, a two-way-coupling is activated to use further result variables. The influence of the particles on the flow is prevented by calculating the steady-state flow solution first without particles. Then the Navier-Stokes equations are deactivated, and one Lagrangian iteration is calculated separately.

In a pre-analysis it was examined whether the evaporation of the water component of the particles has to be considered for the simulation. A mass fraction of about 10.4 % NaCl, and 89.6 % water is assumed [1]. According to the settings of Feng et al. [1] the initial droplet diameters are fixed at 2 μm, which represents the smallest droplet diameter in their study. Compared to the aerosol particle diameter distribution used in this study, this represents a larger particle with a corresponding long evaporation time. The initial mass fraction of 0 % is defined in the domain. The results show that the liquid water only exists for an average time of 0.024 s before it is completely evaporated. During this time, the particle trajectories travel about 1 mm. The range of influence of the gaseous water due to diffusion is limited to about 80 cm.

Due to this small zone of influence and the very short evaporation time of liquid water, the influence of multicomponent particles is neglected in the following. It is assumed that only solid particles are emitted directly from the particle source [17].

### Inert particle setup

The inert particles of solid sodium chloride are injected in a 60 degrees cone shape. The injection occurs in a spatial radius of 0.01 m. The initial velocity of the particles is set to 0.47 m/s at a total flow rate of 8.625·10^−10^ kg/s. The activation of the Saffman Law allows lift forces in shear flows. Based on the measured particle size distribution the analytic Rosin-Rammler distribution is used to adjust the particle diameters in the simulations to the experimental diameter distibution of the aerosol generator. This distribution is defined by the following parameters: minimal diameter 0.19·10^−6^ m, maximum diameter 9.65·10^−6^ m, mean diameter 3.57·10^−6^ m and a spread parameter of 1.90. A stochastically random walk model of 10 leads to a total number of 10,000 trajectories that are calculated in a simulation. The maximum number of steps of 100,000 and a step length factor of 5 defines the tracking parameters and the abort criteria if a particle stream does not reach a target boundary. For the calculation of the trajectories an automatic adaptive time step is used. It is assumed that particles stick on all solid surfaces due to Van der Waals forces [20]. Complete reflection and reentry of particles at the door gap and pressure side of the purifier is assumed. Particles can escape the domain at the intake of the air purifier.

### CFD setup

The turbulence modeling is performed using the *k* – *ε* realizable model with the Menter Lechner wall treatment. It is assumed that the airflow in the room is steady. According to the incompressible ideal gas law, thermally induced buoyancy flows are considered. At low temperatures as well as normal room temperature, heat transfer by radiation can be neglected, [30]. According to DIN EN 13779 [31], persons in the classroom represent a heat source with a heat flux of 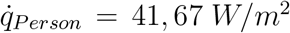. In winter, the window surface temperature cools down and radiators are used [32]. The thermal power loss of the air purifier is adapted according to the power level of the fan. The equations are solved in a coupled scheme. The Navier Stokes equations are discretized with a second order method in space. The two additional transport equations for the turbulence modeling are solved with first order accuracy.

### Case studies

Due to the steady state flow simulation, particle charging and decaying phase can not be considered like in the experiments. Based on a steady state flow field, the Lagrangian solution still provides time-dependent information and allows for a validation by comparison with the experimental results. First, simulations are performed to validate the setup and to compare the results with experimental data. When investigating the influence of the location of air purifier and particle source on local aerosol concentration, the simulations are performed according to the experiments. In addition, the visualization of the room air flow will provide further references for the operating conditions of air purifiers. A comparison of summer and winter case shall determine the influence of cold window surfaces and warm radiators.

## IV). Results and discussion

### IV).1. Experimental results

The experimental results are split up into validation of the measurement method, decay rate comparison of the high-end and low-cost optical particle counters, evaluation of the position and orientation of the air purifier and the influence of the position of the aerosol particle source on local exposure.

#### Validation of instruments and measurement method

In order to validate the measurement method five low-cost OPCs are placed along a grid at face-height of a single thermal dummy and one high-end OPC is placed at table-height for comparison.

**Figure 2:**
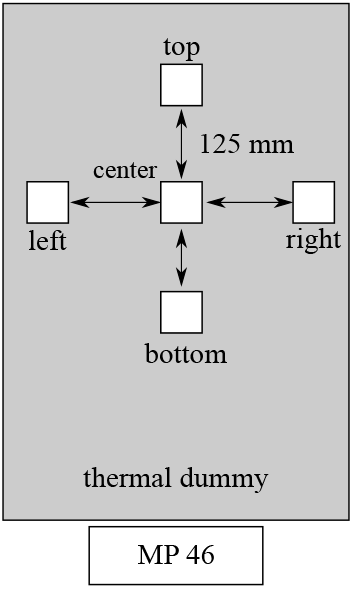
Four low-cost OPCs are cycled around a central OPC at face-height of a thermal dummy. A high-end OPC on the table is used as a reference.

The air purifier is positioned in the south-western corner of the classroom and oriented so that the outlet points towards the front of the room (SW-S). The aerosol particles are emitted at the back of the room near the air purifier (position 80, see Figure 1). After 4 min of measuring the ambient particle concentration, the atomizer (3,5 bar; 16 wt-% NaCl-Solution) and air purifier (ACH 5) are turned on. After 20 min of charging the room with particles, the atomizer is shut off and the air purifier kept running for further 20 min. To distinguish between potential spatial concentration differences, in further experiments the central sensor unit is fixed in place, while the surrounding sensors are cycled around the center point (experiments V1 - V4).

The PM1 exposure determined at each point is set in relation to the center point value. A comparison of the PM1 load at each measuring point (Figure 3 right-hand side) and the PM1 load measured by each device relative to the center point (Figure 3 left-hand side) shows a consistent off-set between the single devices independent of their respective location. The high-end OPC (MP 46) consistently measures a 20 % to 30 % higher PM1 load. This is due to the wider measuring range and a more precise measurement at high particle counts (> 5000 P/cm^3^). Although the exact PM1 mass concentration is subjected to uncertainties due to the varying measurement precision of the low-cost and high-end sensors, the reduction in the relative PM1 exposure at either table height or face height, as well as the decay rate at each measurement point can be used to evaluate the efficiency of the air purifier depending on the operating parameters. The results futher show that a single measurement point at face height is sufficient to identify the PM1 exposure at this location.

**Figure 3:**
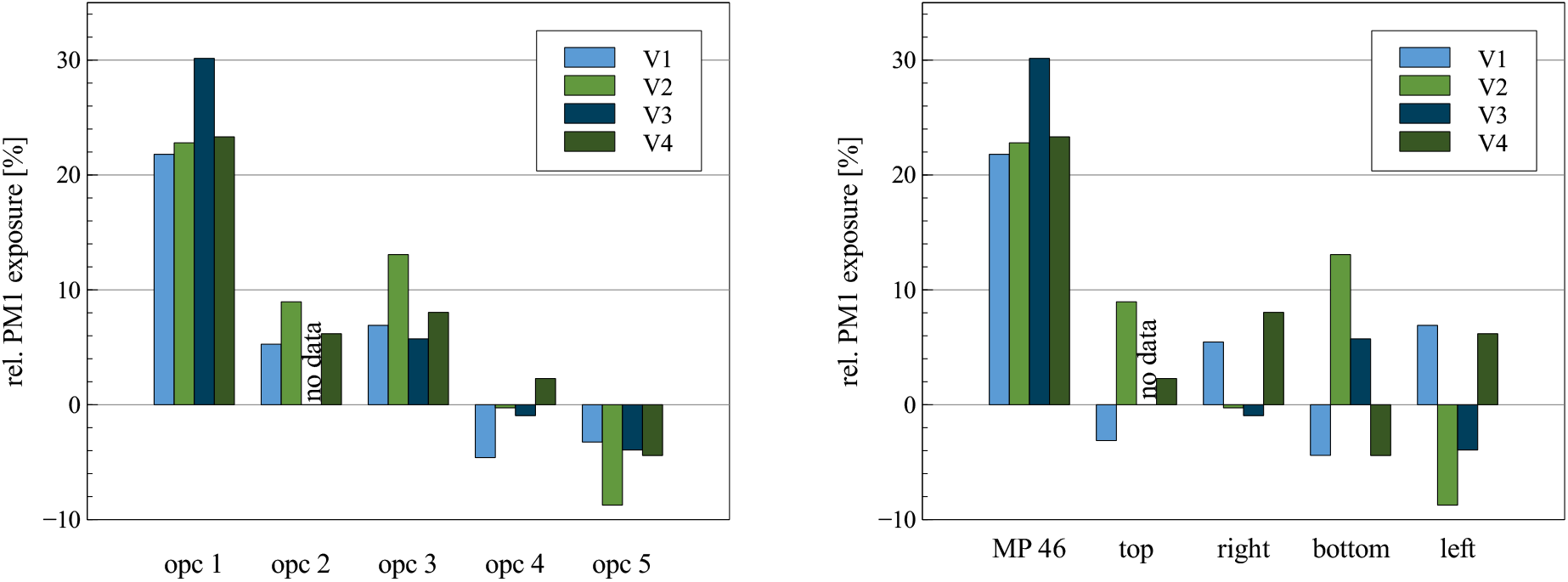
left-hand side: relativ PM1 exposure in relation to the center measurement point by device; right-hand side: relative PM1 exposure measured by location

#### Decay rate of low-cost and high-end sensors

Figure 4 shows the temporal PM1 mass concentration curve of an exemplary validation experiment. Regression coefficients R^2^ > 0.99 show a high correlation between the fitted decay curves and measured data. The decay rate at a single point in the room at an ACH of 5 and 10 are (0.092 ± 0.001) min^−1^ and (0.188 ± 0.002) min^−1^ respectively. In this case 50 % of the initial particle concentration decay after (7, 52 ± 0, 09) min at an ACH of 5. Increasing the ACH to 10 gives a half-life time of (3, 63 ± 0, 04) min, while 99% of the initial particle concentration decays after (24.6 ± 0.7) min.

**Figure 4:**
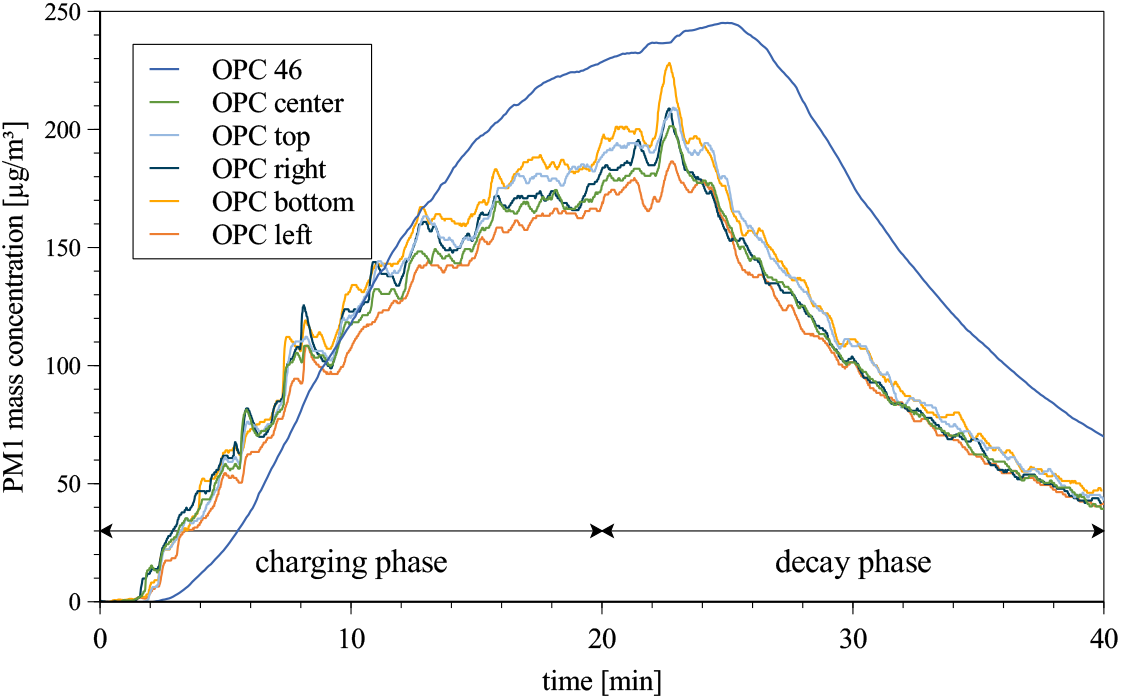
Temporal PM1 mass concentration of an examplary valdidation experiment showing the charging and decay phase

##### IV).1.1. Evaluation of the position and orientation of the air purifier

The influence of the position and orientation of the air purifier on distribution and decay of the aerosol particles was evaluated by comparing the PM1 exposure at 4 measurement points using the high-end OPCs. In Figure 5 (left-hand side) the PM1 exposure at each measurement point is shown. The location of the air purifier has a noticable impact on the distribution of the emitted particles. While in absence of an air purifier, the emitted particles initially follow the thermally induced air flow by the measurement point MP 58 (located between the particle source in the middle of the room and the window front), operating the air purifier leads to increased PM1 concentration at measurement points close to the intake. For the positions NE and N this is MP 58. Position NW and SW show increased concentration at points MP 86 (back of the room) and MP 52 (near wall). In case of NE-E the emitted particles travelled to the intake without passing by the OPC, leading to a minimal PM1 exposure measured in this set-up. This example clarifies that a high spatial resolution is necessary especially during the charging phase where it can not be assumed that the particle concentration is homogeneous throught the room.

**Figure 5:**
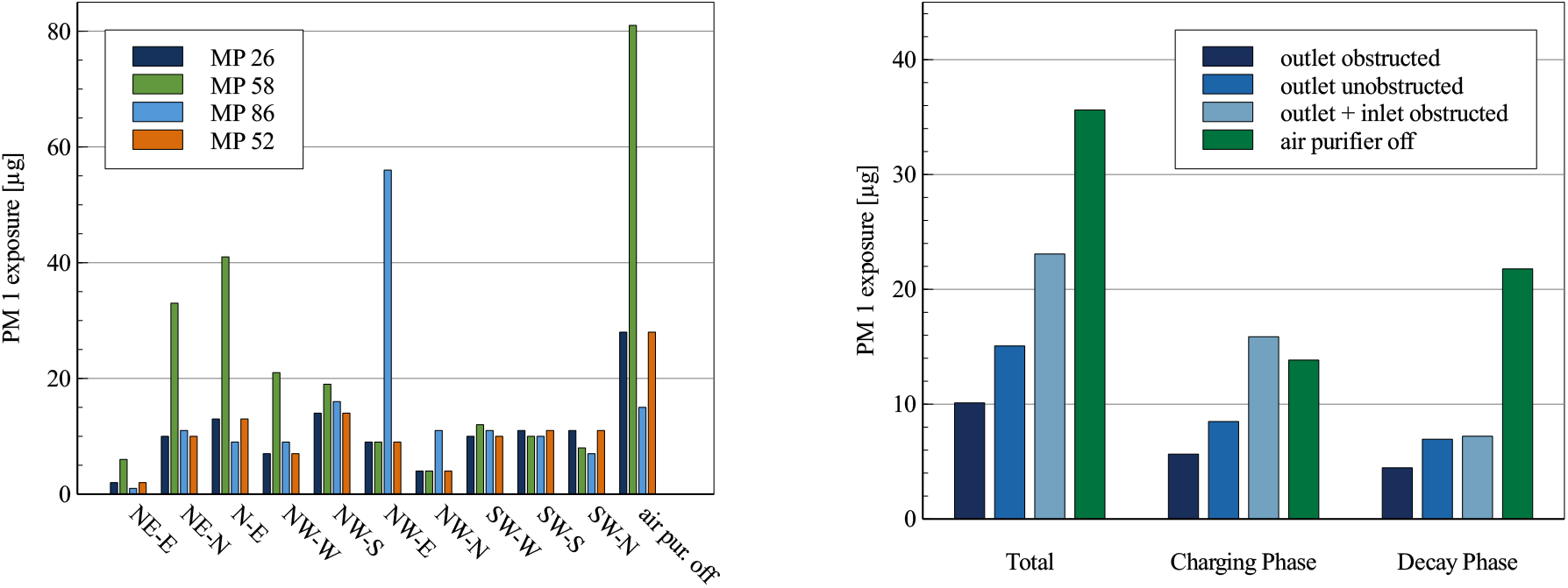
left-hand side: PM1 exposure at each measurement point for all investigated set-up parameters at ACH 5; right-hand side: averaged PM1 exposure by set-up catagory at ACH 5

In order to evaluate the set-up parameters of the air purifier, the measured PM1 exposures where averaged within three set-up categories: I. the air purifiers outlet is obstructed (oriented in such a way that the outlet points towards the wall or window), II. the outlet is unobstructed (oriented in such way that the outlet points freely into the room), III. outlet and intake are obstructed (oriented in such a way that both the outlet and the intake are obstructed by either a wall or a window).

Figure 5 (right-hand side) shows the average PM1 mass load for category I, II and III over both charging and decay phase as well as split up into each phase.

In all cases, the air purifier reduces the particle load compared to the natural decay. Setting up the air purifier by obstructing the outlet leads to an average 70 %reduction of the total PM1 exposure over the charging and decay phase. By pointing the outlet stream unobstructed into the room, the PM1 load is only reduced by an average of 58 %. This is attributed to the outlet stream distributing the emitted aerosol particles throughout the room during the charging phase rather than depositing them. Obstructing the outlet as well as the intake of the air purifier is particularly disadvantageous, leading to an average PM1 load reduction of only 30 %, while showing a PM1 exposure comparable to the absence of the air purifier during the charging phase. It can be concluded that orientation of the air purifier has a significant impact on the efficiency of the reduction of the airborne particulate matter. In this case, obstructing the outlet stream reduces the distribtion of the aerosol particles throughout the room. This ensures that the air purifier is employed at maximum efficiency.

##### IV).1.2. Influence of aerosol particle source on local exposure

Derived from the results of the first experimental set-up, four case are investigated further (SW-W, SW-S, SW-N, no air purifier). The new experimental set-up includes 9 thermal dummies equipped with low-cost sensors at face-height as well as varying the position of the aerosol particle source. The PM1 exposure over the course of the experiment is set in relation to PM1 load measured using no air purifier (see Figure 7 left-hand side).

**Figure 6:**
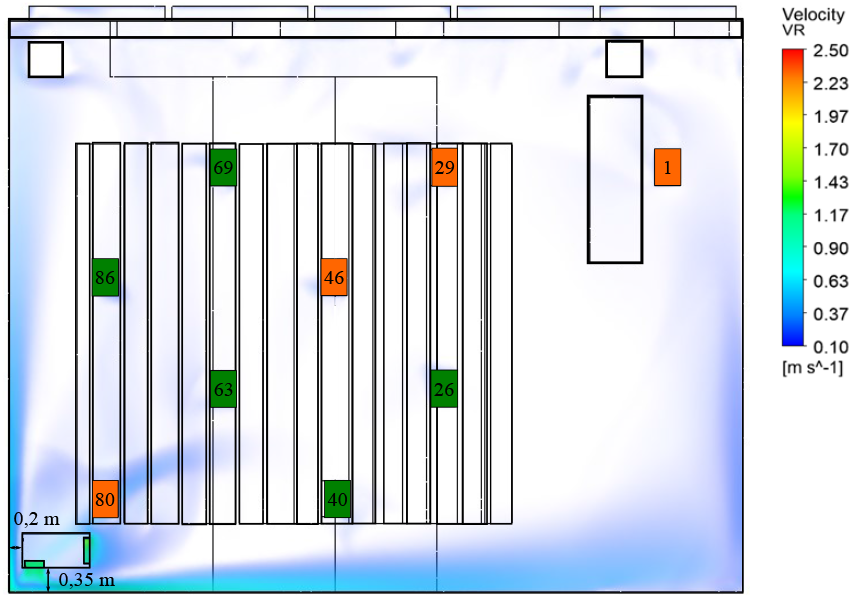
Experimental set-up showing location of the air purifier, the aerosol source positions, measurement points and air velocity larger than 0.1 m/s for position SW-W at ACH 5

**Figure 7:**
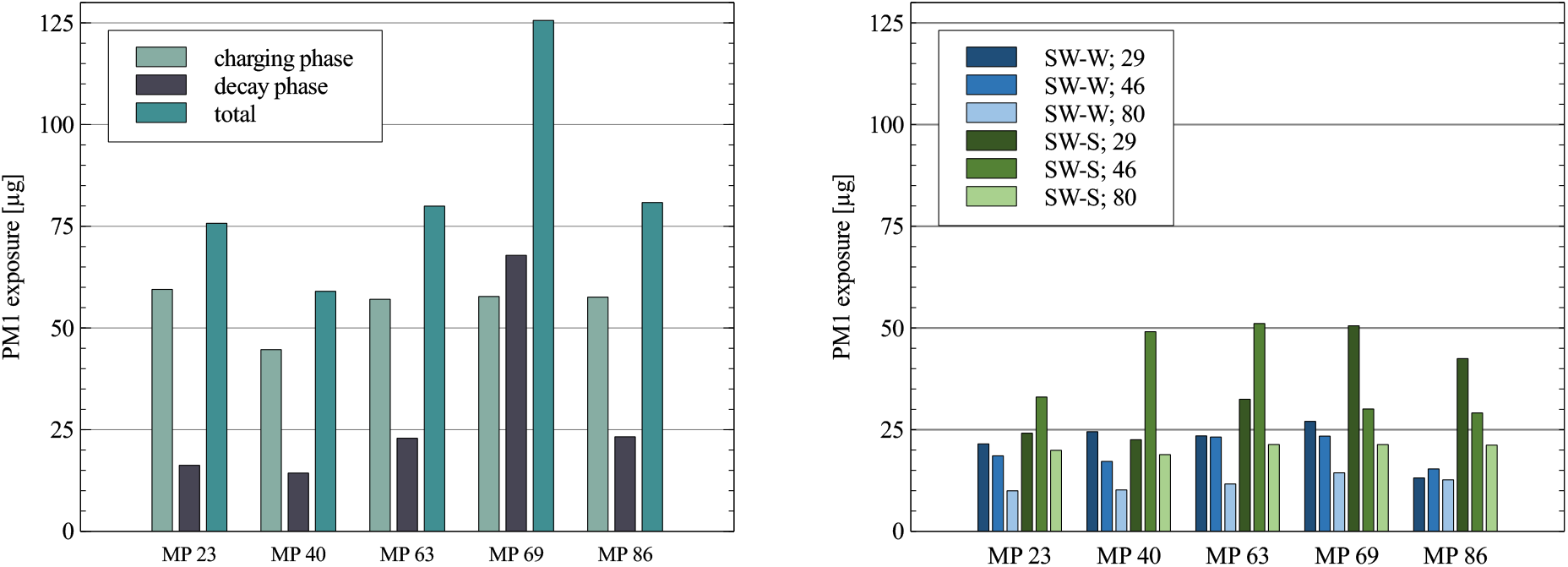
left-hand side: PM1 exposure at face height in the absence of the air purifier; right-hand side: PM1 exposure at face height for the positions SW-W and SW-S depending on the aerosol source (29,46,80) at an ACH of 5

Overall the use of the air purifier decreases the PM1 load independently of the aerosol source compared to no air purifier (see Figure 7 right-hand side). The highest reduction was achieved with the set-up parameter SW-W (at ACH 5), decreasing the PM1 exposure throughout the room on average by 75% independently of the aerosol source position. This is slightly higher than the reduction determined in the first experimental set-up, due to the fact that emitting particles near the intake leads to an overall reduction of well above 80%. SW-N decreases the PM1 exposure throughout the room on average by 61 %. In this worst-case scenario the aerosol particles are emitted from seat 29 at a significant distance to the air purifier. Further dependency on the aerosol source position is not investigated. The last case with an unobstructed intake and outlet (SW-S) decreases the PM1 exposure throughout the room on average by 61 %. This is very close to the 58% reduction of PM1 particle concentration measured in the first experimental set-up for unobstructed intake and outlet. Regarding the charging phase, SW-S leads to localized increases of PM1 exposure compared to no air purifier, further indicating, that an unobstructed outlet negatively effects the removal of the aerosol particles. SW-W holds up to be the best set-up case, showing the lowest increase in PM1 exposure during the charging phase independently of the location of the aerosol source and indicating that obstructing the outlet air flow is a viable strategy in preventing the distribution of particles throughout the room.

### IV).2. Numerical results

#### Validation of the numerical model

The impact of the mesh on the results of the CFD simulation is investigated. Starting from a fine mesh, the grid is coarsened and the resulting differences are evaluated.

**Figure 8:**
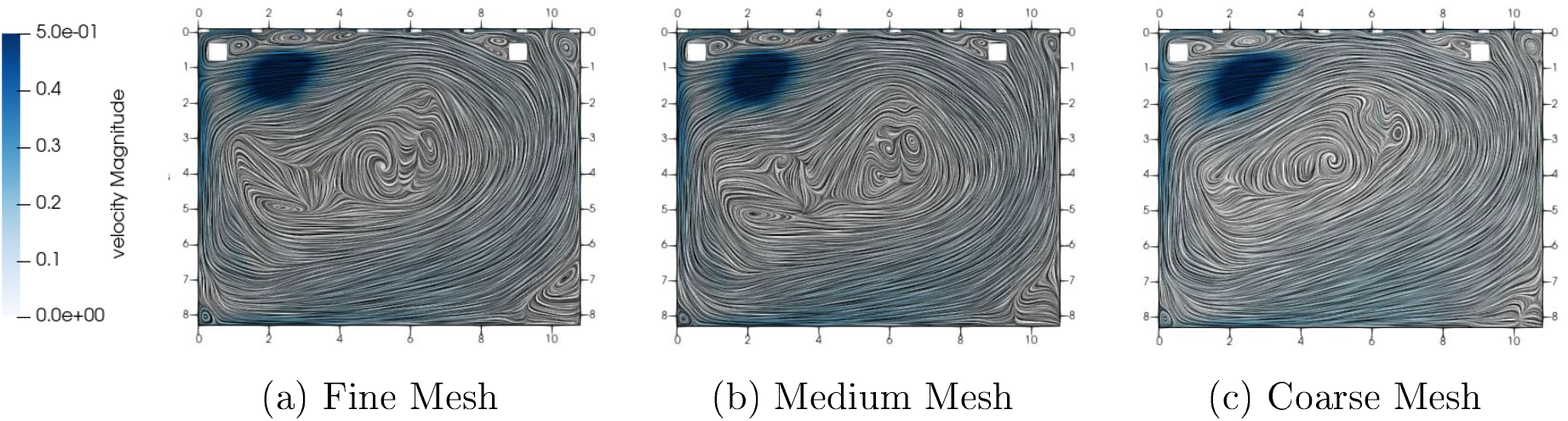
Comparison of vortex structures in slow flow regions on different meshes

Using fine and medium mesh similar vortex structures can be seen at head height of 1.7 m. In contrast when using the coarse mesh, different flow vortices occur in the center of the room. Even though the faster flow velocity larger than 0.1 m/s of all mesh variants corresponds very well with the results using the fine mesh, different aerosol particle dispersion occurs. The particle trajectories take different paths because of the slow flow vortices in the center of the room, where the particles are injected. The medium mesh settings are used as a compromise between computational duration and accuracy.

#### Validation using steady particle tracking

A validation of the simulation results is done based on the stationary particle simulation, since this is also used in the simulation study. Here, temporal information is only available in the Lagrangian phase.

After 60 s, the particles have barely dispersed. No measuring device should react to the turned-on aerosol source. After about two minutes, the particles have arrived at MP58 and MP86. The MP58 is exposed to the most particles. The MP26 and MP52 do not show any measurement data yet. After about three minutes, the particle trajectories pass the MP52. At MP26 near the placement of the air purifier the first particles from the aerosol generator are measured only after four minutes. This matches to the experimental data in Figure 10.

**Figure 9:**
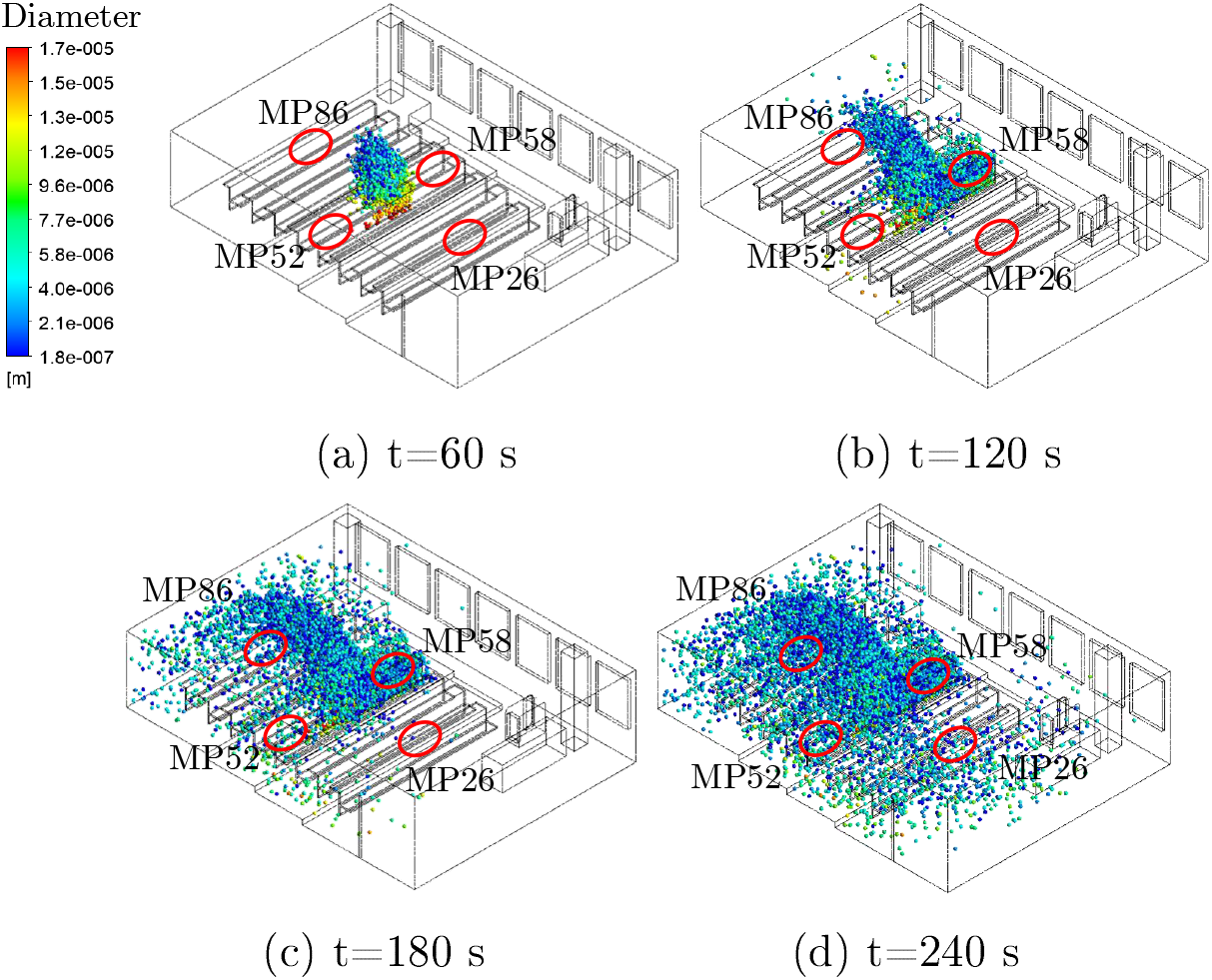
Dispersion of Lagrange particles using steady particle tracking for validation with measurement data

**Figure 10:**
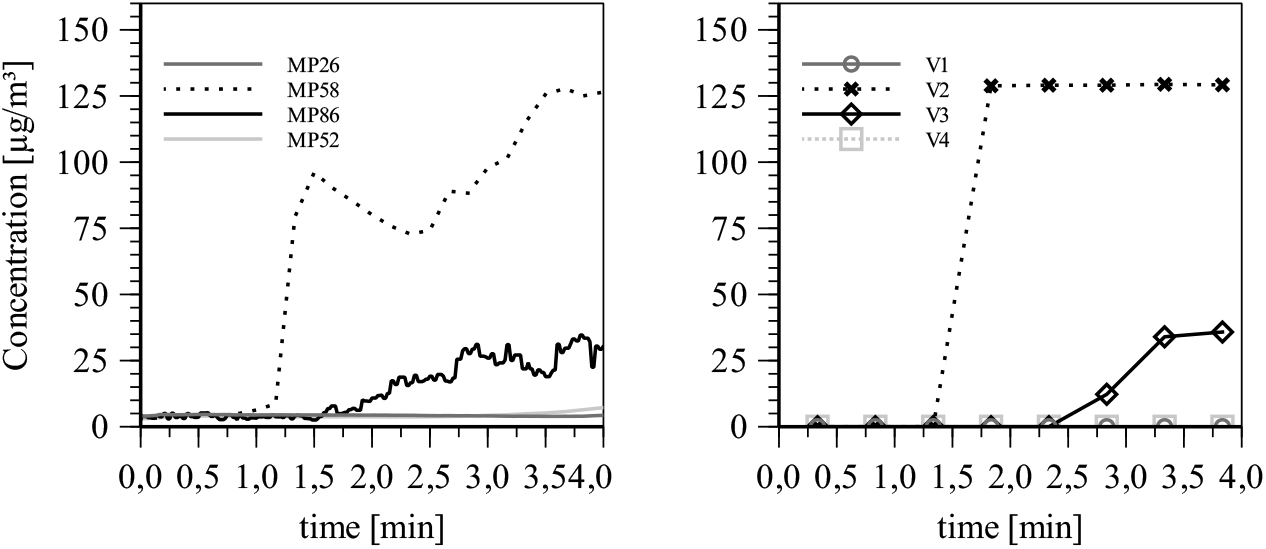
Comparison of particle mass concentration determined experimentally (left-hand side) and unsteady simulation (right-hand side) at four different measurement locations at ACH of 5

#### Validation using unsteady particle tracking

A transient particle simulation can offer further information for the validation of the simulation settings to the experimental measurement results. To evaluate the mass concentration spatially and temporally, control volumes have to be defined according to the position of the OPCs.

The order of magnitude, as well as the time evolution of the mass concentration at the control volumes matches to the experimental data at the measurement locations.

#### Influence of air purifier and particle source on local particle exposure

The analysis of the air flow at different positions of the air purifier already gives an insight into the aerosol distribution. According to DIN EN ISO 7730 a maximum average air velocity of 0.24 m/s is specified for classrooms [33]. In all cases, the flow velocity at the occupied seats is low enough not to cause thermal discomfort.

**Figure 11:**
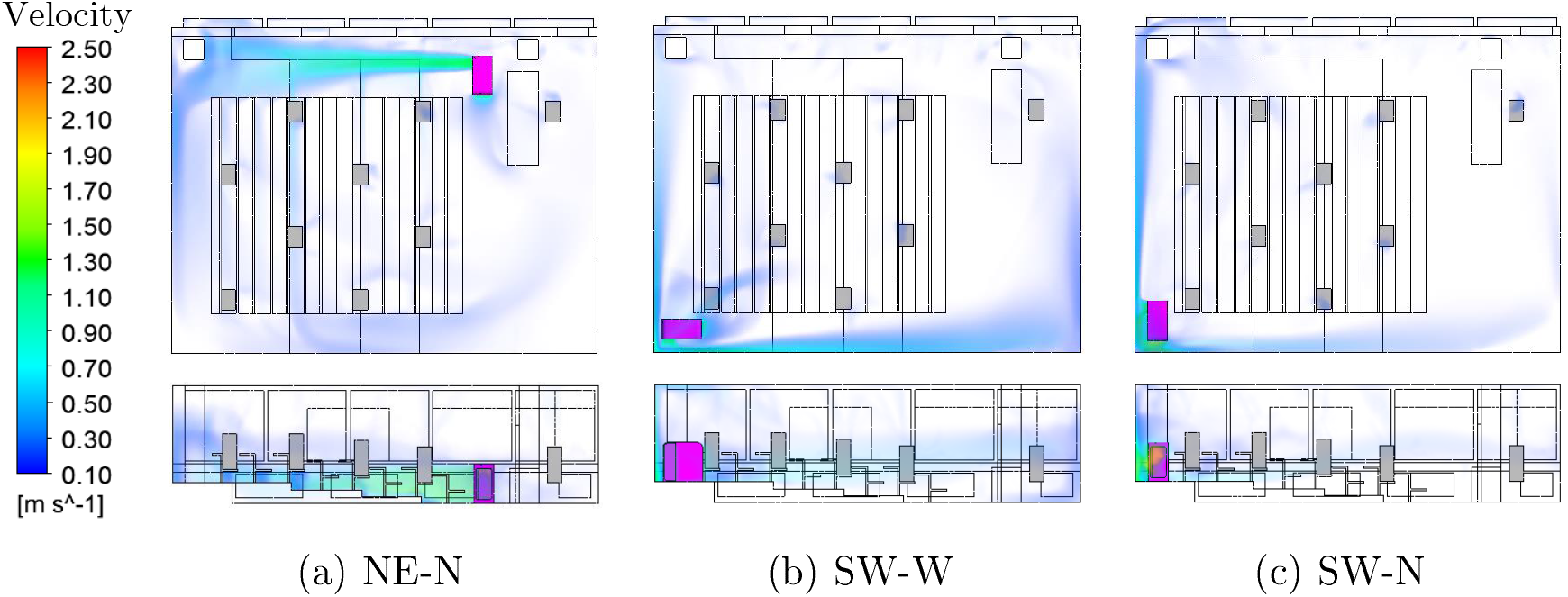
Comparision of different air purifier positions and air velocities at ACH of 5

The fates describe the number of particle streams arriving at a defined target boundary. The different labels represent the different particle source positions. The data series represent the different locations of the air purifier in Figure 12. Particles that reach the air purifier can be removed from the ambient air. This reduces the amount of potentially infectious particles. The plotted data shows the relative number of particle streams that are collected in the filter at an ACH of 5, allowing to compare different positions and orientations of the air purifier at the same filter volume flow.

**Figure 12:**
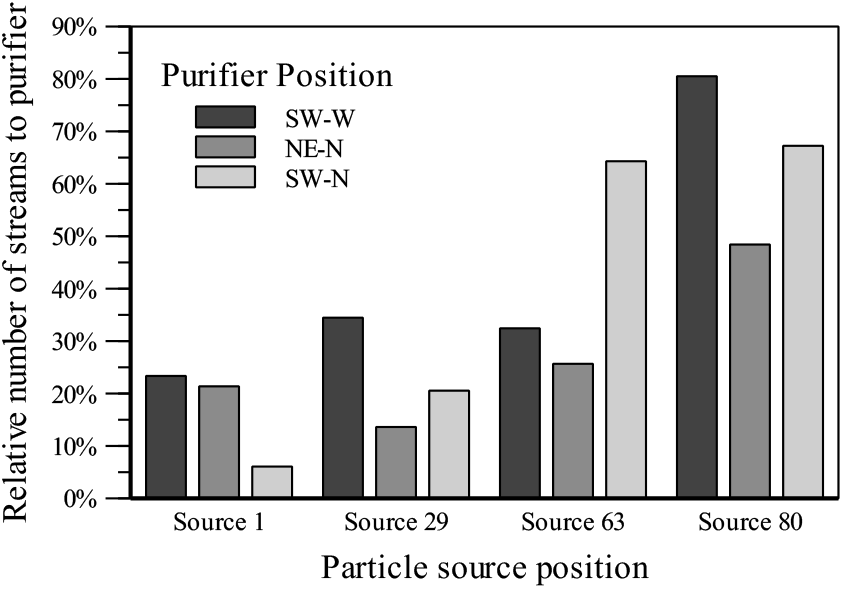
Particle fates for different purifier positions and different particle source positions at an ACH of 5

In air purifier position SW-N, it is noticeable how strongly the number of filtered streams depends on the source position. In the worst case, only about 5 % of the particle streams reach the filter. Due to this strong dependence of the source position, the purifier position SW-N is not recommended. The position NE-N shows the best uniformity of the number of filtered streams. However, this is at a generally low level, so that on average only about 28 % of the injected particle streams arrive at the air purifier. Position SW-W achieved the best results in this comparison. On average, about 40 % of the emitted particle streams reach the room air purifier. In the worst case, still about 25 % of the particle streams are removed by the filter. The positioning of the room air filter with outflow against the wall and intake in the direction of the room interior shows the best results in this simulation comparison. The visualization of the mass concentration allows qualitative evaluations of the particle paths from the injection to their target boundary. A column represents one particle source position and allows for a comparison of the different locations of the air purifier in different rows in Figure 13.

**Figure 13:**
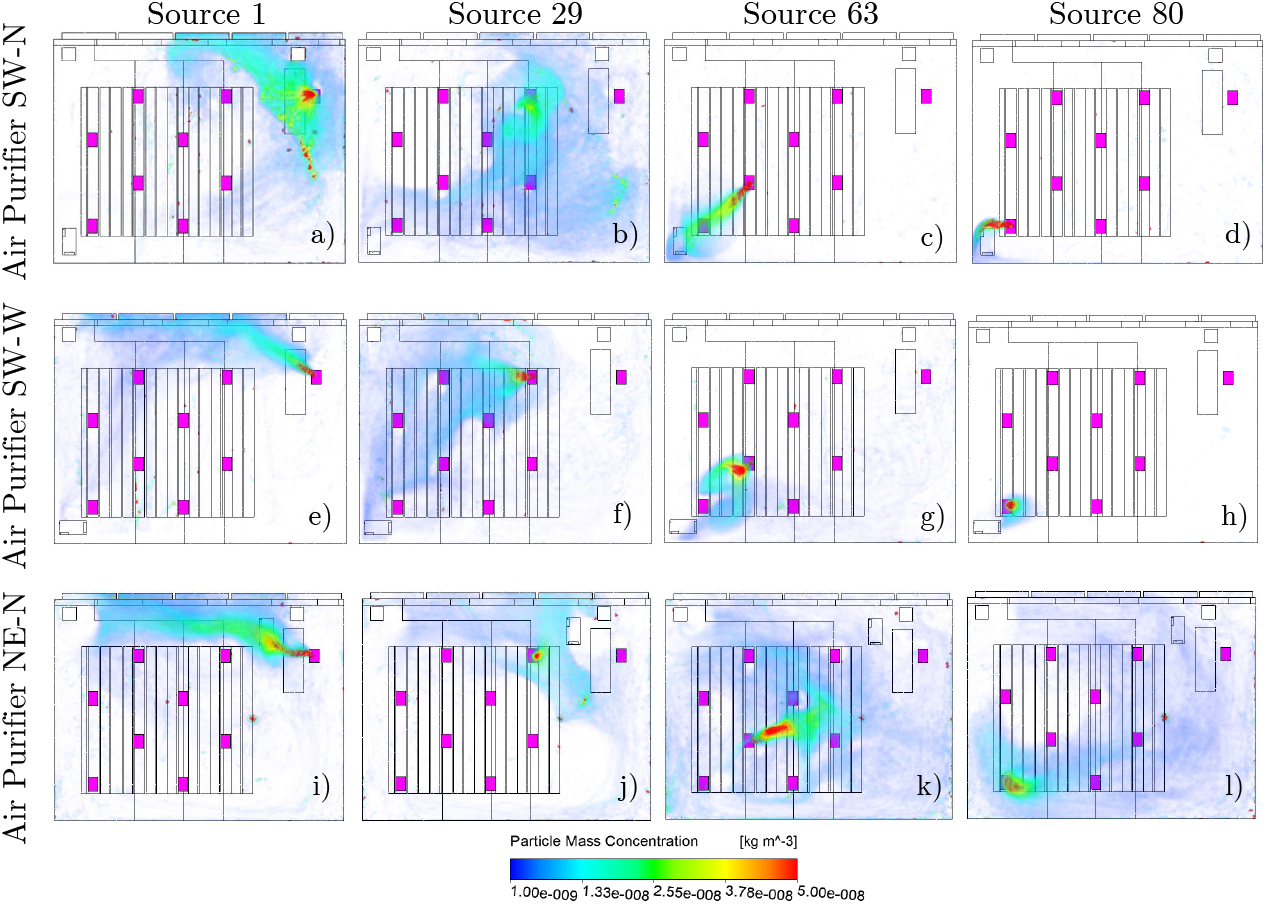
Particle concentration of different air purifier positions and paricle source positions at an ACH of 5

#### Extended cases studies

As an extension of the numerical study, the flow velocity at different ACHs and the differences in buoyancy flows in summer and winter will be investigated.

#### Air velocity for different ACHs

The zone of influence of the intake of the air purifier is significantly smaller than the pressure side area. This value is exceeded at the persons seats when the air purifier is operating at the highest level. At an ACH of 5 these comfort limits are just fine at the seating areas. This level represents the maximum performance of the room air filter in this room and this positioning without causing discomfort to the persons due to excessive air velocities. Operation of the room air filter at ACH of 5 is recommended in this room to achieve the best possible compromise between comfort due to air velocity and filter volume flow. The air purifier has to be oriented in such a way, that the outflow area is not directed towards occupied seats. In addition, by variation of the ACH, it can be shown that the duration of the particle movement from the source to the air purifier decreases with higher flow rate. Depending on the air change rate, redirecting particles to the air purifier reduce the possible infectuous particle streams by more than 50 %.

**Figure 14:**
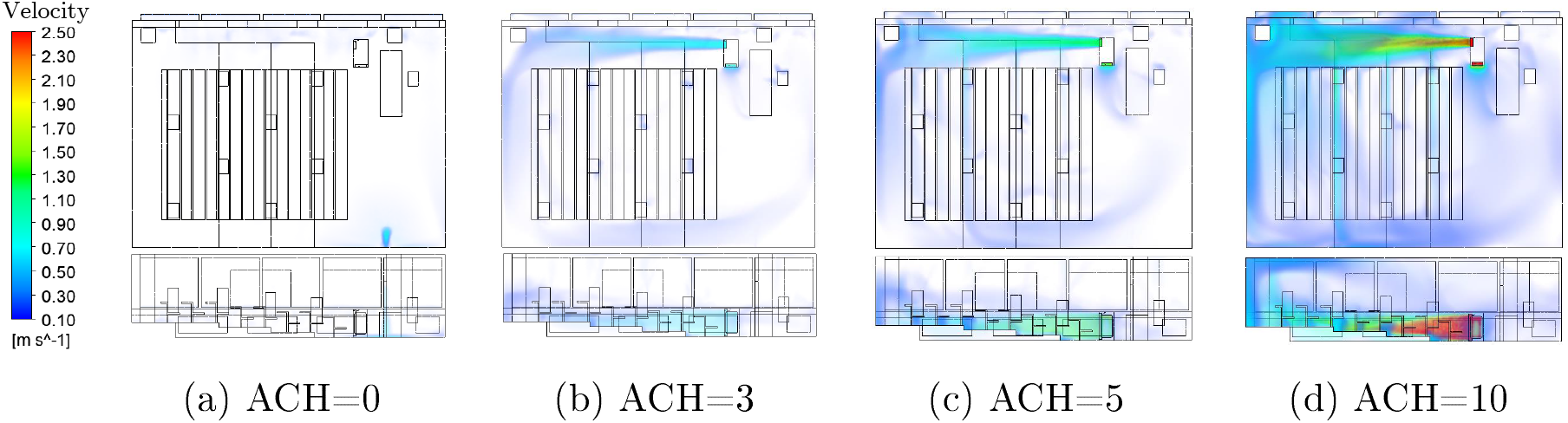
Air velocity (NE-N) at different ACHs

**Figure 15:**
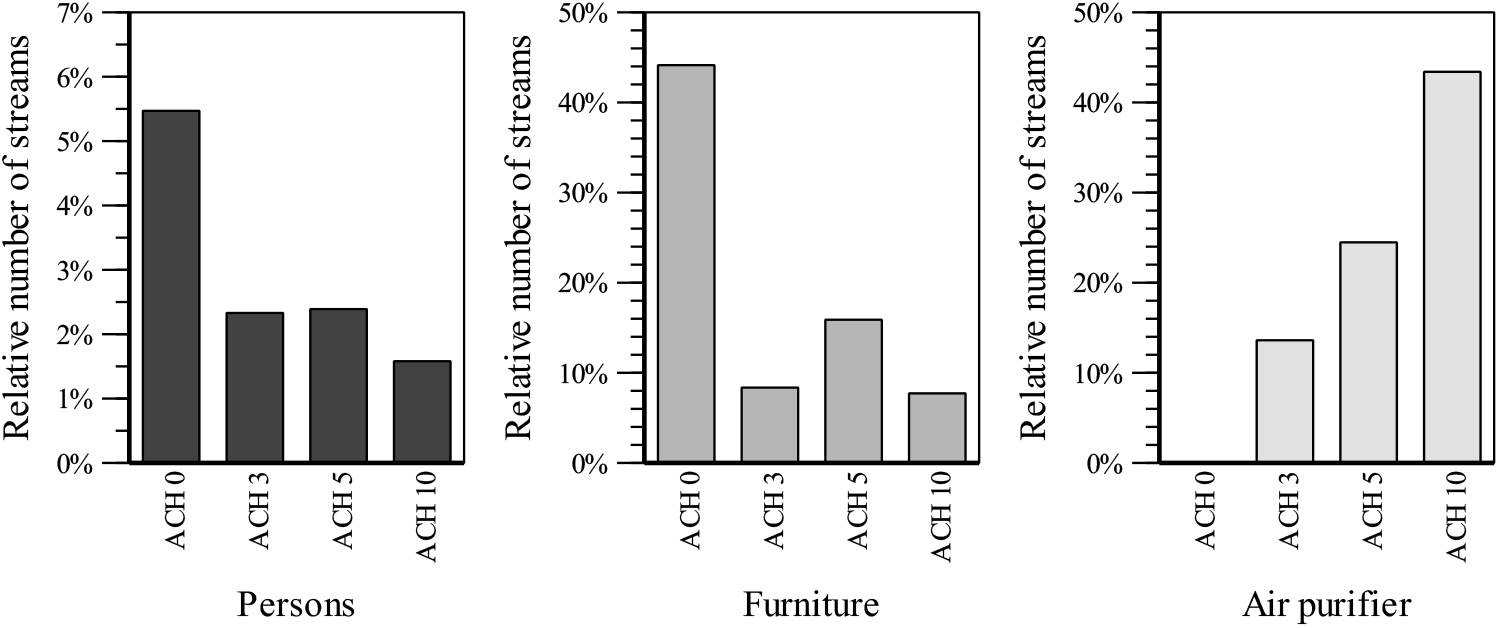
Comparison of particle fates at different ACHs

#### Summer vs. winter air flow

Using the incompressible ideal gas law, the air density can change due to local temperature differences. The density differences, lead to buoyancy forces. In Figure 16, the vertical velocity (v in positive Y-direction) is shown.

**Figure 16:**
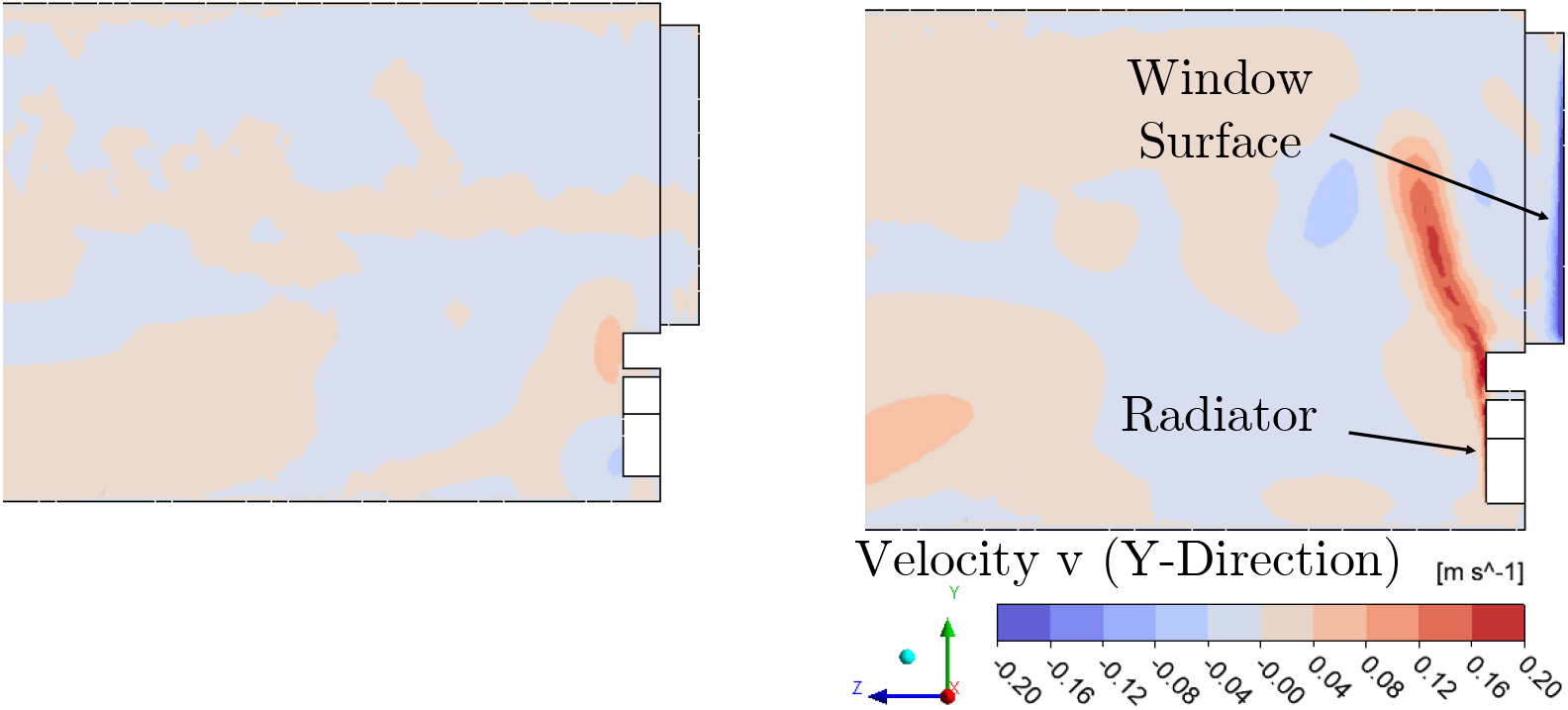
Comparison of summer (left-hand side) and winter (right-hand side) air flow near windows and radiator

The difference of summer and winter simulation is mainly noticeable at the window side. The cold window glass in winter leads to sinking air flows. Directly below the window is the radiator, which causes upward flows. At the height of the window sill, these descending and ascending streams intersect and lead to a lateral deflection of the heating flow towards the interior of the room. However, the area of influence of the heating flow is locally limited to about one meter.

### IV).3. Limitations of the presented approach

While the PM1 exposure is used to evaluate the efficiency of the air purifier, it is unlike the inhaled particulate matter responsible for airborne transmission. First, this is due to inhaled particles being deposited in the lung and therefore removed from further circulation. Secondly, the actual exposure to virus-laden droplets is far lower than the exposure measured in this study. For this reason, no direct infection risk was calculated, but rather the relative infection risk connected to the optimized operating parameters of the air purifier evaluated. In a real-world application, the HVAC system is likely to be running, influencing the distribution and decay of particle concentration. Furthermore, the examined cases did not include dynamic processes such as moving particle sources and varying source strength (through coughing, talking etc.). Since most air purifiers on the market today are highly distinguishable in their design regarding the location of the intake and outlet, this is a highly specific case from which no generalization of the observed phenomenon to all types of air purifiers can be made. Although the low-cost OPCs have a much lower resolution, as well as a smaller measurement range compared to the high-end OPCs, the validation experiments showed that the decay rate are highly comparable to the latter devices. Since the high-end OPCs measure considerably more PM1 mass concentration over the course of the experiment, the efficiency of the air purifier was evaluated either by comparing the low-cost OPC or high-end OPC data. In order to push the particle size distribution into the measurement range of the low-cost OPCs, the mass concentration of the saline solution was raised from 2,5 wt-% in the first experimental set-up to 16 wt-% in the second. This raised the median particle diameter *x*_50,0_ from 0.25 μm to 0.34 μm. The experiments were carried out over a substantial period in winter going into spring. The ambient conditions changed accordingly and influenced parameters such as outside temperature and relative humidity, which will influence the thermal convection of particles exposure throughout the room. The most critical point is the assumption that particle fates correlate with the infection risk. This correlation has not been proven and is an assumption. In addition, it is assumed that each stream has the same potential for infection. Obviously, not only the destination of the particle streams is relevant for the infection risk. Particles can be infectious on their path between injection and destination boundary. For this reason, the particle mass concentration was calculated. From the experimental results it is known that the particle mass concentration with build-up phase and decay phase is strongly time-dependent. The numerical result is only available as stationary averaged values in time. Therefore, it can only be used as a subjective evaluation criterion. A limited number of 10,000 trajectories are calculated in the simulation, although in reality the number of particles is essentially higher. Another aspect are differences of the numerical model from reality. First, the geometric model is highly simplified. In addition, the real boundary conditions of the room are difficult to detect and to transfer into the simulation boundary conditions.

## V). Conclusion

In this work, we have studied the influence of position and orientation of an air purifier on the aerosol particle concentration in a lecture hall and identified preferable cases with respect to a small aerosol exposure of persons present in the room. Specifically, we have set up a PM1 sensor network consisting of 12 sensors exactly at the locations where particles may be inhaled and included the influence of thermal buoyancy by mimicking the effect of persons present in the room using thermal dummies. We investigated both the charging phase where the air purifier fights against a source (“infected person is in the room”) and by studying the decay phase (“infected person has left the room”) while using an unrealistically high emission rate to cut the experiment short. Furthermore, we validated a CFD model based on the measurements in order to study particle fates and to allow for the investigation of situations, which can hardly or not be considered experimentally.

Overall, the results stress that filtration is an effective means of reducing aerosol particle concentrations. The measurements suggest that a blowout against the wall may be particularly advantageous in order to avoid an increase of local particle concentration at the locations where persons breathe after turning on the air purifier. In turns out that the air purifier can very effectively reduce aerosol particle concentrations in a combined loading and decay scenario by 86 % using a good orientation with obstructed outlet and by 61 % in an unfavorable orientation and position. The CFD simulations suggest that an additional effect of risk mitigation is the fact that the air purifier reduces the deposition of aerosol particles on critical surfaces (persons, furniture). However, it needs to be stressed that aerosol exposure and particle fates may differ from the actual infection risks due to system-inherent differences such as the fact that inhaled particles are removed in reality, but remain within the system in the models and in the experiments.

Further work needs to be done to provide insight into the fact how the specific flow configuration influences the results. Therefore, this study should be extended taking into account different air purifier devices. In addition, it needs to be investigated to which extend the results obtained depend on the specific geometry of the lecture hall and in how far they can be generalized.

## List of Figures

1. left-hand side: position of the air purifier, the receiving and emitting thermal dummy (marked by green and orange circles) and the high-end OPCs(marked blue); right-hand side: picture of the lecture hall with thermal dummies and sensor network ...... 7

2. Four low-cost OPCs are cycled around a central OPC at face-height of a thermal dummy. A high-end OPC on the table is used as a reference ......... 14

3. left-hand side: relativ PM1 exposure in relation to the center measurement point by device; right-hand side: relative PM1 exposure measured by location .... 15

4. Temporal PM1 mass concentration of an examplary valdidation experiment showing the charging and decay phase .... 16

5. left-hand side: PM1 exposure at each measurement point for all investigated set-up parameters at ACH 5; right-hand side: averaged PM1 exposure by set-up catagory at ACH 5 .... 17

7. left-hand side: PM1 exposure at face height in the absence of the air purifier; right-hand side: PM1 exposure at face height for the positions SW-W and SW-S depending on the aerosol source (29,46,80) at an ACH of 5 ..... 19

8. Comparison of vortex structures in slow flow regions on different meshes ... 20

9. Dispersion of Lagrange particles using steady particle tracking for validation with measurement data .... 21

10. Comparison of particle mass concentration determined experimentally (left-hand side) and unsteady simulation (right-hand side) at four different measurement locations at ACH of 5 .... 21

11. Comparision of different air purifier positions and air velocities at ACH of 5.... 22

12. Particle fates for different purifier positions and different particle source positions at an ACH of 5 .... 22

13. Particle concentration of different air purifier positions and paricle source positions at an ACH of 5 .... 23

14. Air velocity (NE-N) at different ACHs ..... 24

15. Comparison of particle fates at different ACHs ..... 24

## Statements

The data that support the findings of this study are available from the corresponding author upon reasonable request. This research was partly funded by the Ministery of Science, Research and The Arts (MWK) of the State of Baden-Württemberg in the frame of the special funding track COVID-19 research, part of the measures for fighting the SARS-CoV-2 pandemic in the frame of medical research. The authors declare no conflict of interest. The manuscript does not contain experiments using humans or animals. A patient consent statement is not relevant. The authors have not used materials from other sources.

## Acknowledgements

This research was partly funded by the Ministery of Science, Research and The Arts (MWK) of the State of Baden-Württemberg in the frame of the special funding track COVID-19 research, part of the measures for fighting the SARS-CoV-2 pandemic in the frame of medical research.

